# Consequences of a short-term exposure to a sub lethal concentration of CdO nanoparticles on key life history traits in the fruit fly (*Drosophila melanogaster*)

**DOI:** 10.1101/2020.07.15.204370

**Authors:** Samar El Kholy, John P. Giesy, Yahya Al Naggar

**Author notes:** Corresponding author: Yahya Al Naggar, Ph.D., General Zoology, Institute for Biology, Martin Luther University Halle-Wittenberg, Hoher weg 8, 06120 Halle (Saale), Germany.

## Abstract

Nanoparticles of cadmium oxide (CdO NPs) are among the most common industrial metal oxide nanoparticles. Early adulthood (F_0_) fruit flies (*D. melanogaster*) were exposed for 7 days to a sub lethal concentration (0.03 mg CdO NPs/ml, which was 20% of the LC_50_), spiked into food media to test for long term-effects over time and beyond their direct exposure on key life history traits. Effects on survival, developmental time, eclosion rate, fecundity and negative geotaxis performance were assessed. Potential effects on ultrastructure of mid gut cells were also investigated by use of electron microscopy. All studied life history traits, as well as climbing behavior were adversely affected by exposure to CdO NPs. In non-exposed progeny (F_1_) of adult flies (F_0_), a blistered wing phenotype was also observed. Lysis of nuclear and rough endoplasmic reticulum (rER) membranes, mitochondrial swelling and lysis were among the most common cellular alterations observed in midgut cells of F_0_ flies exposed to CdO NPs. Genes encoding for metallothionein (MTn A-D) were significantly upregulated in both parent flies (F_0_) and their progeny (F_1_) after exposure of F_0_ flies to CdO NPs, compared to unexposed, control flies, a result which indicated potential, long-term effects. Taken together, these results suggest that short-term exposure to a sublethal concentration of CdO NPs is sufficient to cause long-lasting, harmful effects on fruit flies.

**Graphical Abstract:** 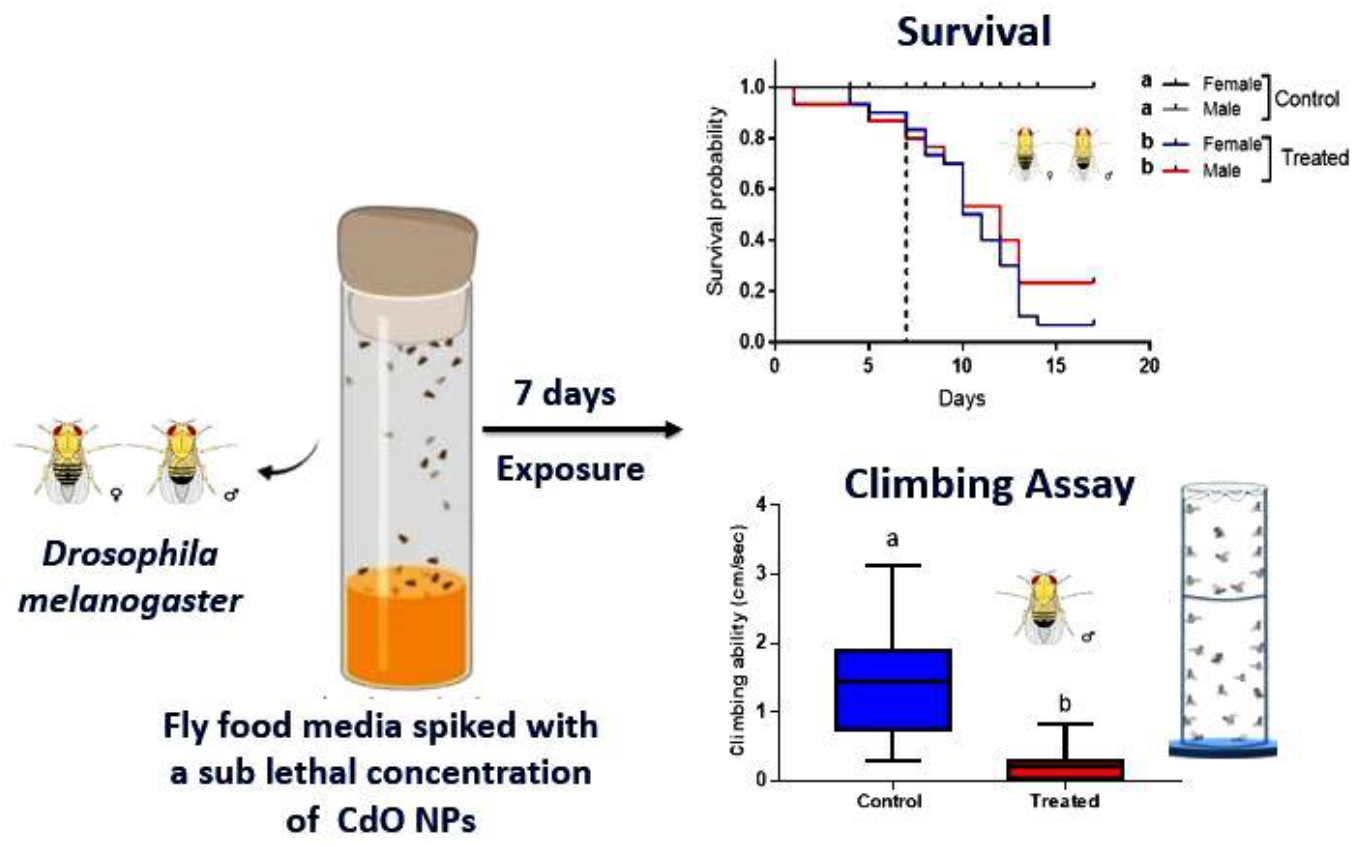

## 1. Introduction

Among several human activities, including engineering, agriculture, manufacturing, medicine and public health, nanotechnology has gained considerable public attention. Since nanomaterials are part of our everyday lives, exposure of humans and wildlife to nanomaterials is inevitable and as a result, research into possible effects of nanotoxicity is increasing (Jeevanandam et al., 2018; Ray et al., 2009). The specific properties of nanoparticles (NPs) allow them to enter organisms and to be transported to tissues, cells, and even organelles in such a way that larger particles may not (Sukhanova et al., 2018). This has raised possible threats to human health and the environment (Hegde et al., 2015; Maurer-Jones et al., 2013; Ray et al., 2009).

Cadmium (Cd) is an inner transition metal occurring naturally in the atmosphere and as an industrial and agricultural pollutant. It has a significant toxic potency of ongoing concern because its concentrations in the environment have increased due to continued mobilization and release by activities of humans. It has various toxic effects, including an extraordinarily long biological half-life of about 20–30 years in humans, low body excretion rate, and storage primarily in soft tissues, mainly liver and kidneys (Naggar et al., 2014; Rafati Rahimzadeh et al., 2017; Rani et al., 2014). Therefore, more attention should be paid to possible toxicity of Cd (Nordløkken et al., 2015; Ye et al., 2015).

Nano-Cd oxide (CdO) is the initial substrate for the manufacturing of quantum dots (QDs) in both medical diagnostic imaging and controlled therapy (Bentolila et al., 2005) and in solar cells (Mane et al., 2006). It is present in the air in industrialized countries, including cigarette smoke (Dumkova et al., 2016a), sediments of lakes and streams (Rzętała, 2016), fertilizers and waste sludge (Davis, 1986). Releases of Cd NPs into the environment could result in their accumulation in the food chain and increased human exposure that could potentially affect health of humans, biodiversity, and the environment.

Several studies have demonstrated toxic potencies of cadmium-containing nanoparticles (NPs) *in vitro* and *in vivo* (AL Naggar et al., 2018; Brunetti et al., 2013; Dabour et al., 2019; Demir et al., 2020) and on human health (Blum et al., 2012; Dumkova et al., 2016b). Although these studies provide independent empirical support for the toxicity of these Cd-containing NPs, however more information about potential risks of these metallic nanoparticles on living organisms are still required; such as for example their effects on life history traits, possible short or long term effects on physiology at the genomic and proteomic levels, during and after exposure.

Here, the fruit fly (*Drosophila melanogaster*) was used as an *in vivo* model organism. Employing *D*. *melanogaster* is less problematic in ethical terms than using other *in vivo* higher animal models. Moreover, around 50 percent of proteins and about 75 percent of genes of human disease exhibit associated sequences in *D. Melanogaster*, meaning that findings obtained for the fly are important for predicting possible effects in other species (Lloyd and Taylor, 2010; Reiter, 2001; Velentzas et al., 2015). In a series of experiments, adult flies were used to test long-term consequences of short-term sublethal exposure of CdO NPs early in life on survival, developmental time, fecundity, climbing activity and the expressions of selected detoxification encoding genes. Phenotypic and ultrastructural effects were also been investigated. We therefore quantified whether short episodes of CdO NPs exposures early in adult life have long lasting effects on life history traits such as fecundity well beyond exposure times.

## 2. Material and methods

### 2.1 Preparation and Characterization of CdO NPs

Cadmium oxide nanoparticles (CdO) were prepared according to methods previously published (DurgaVijaykarthik, D; Kirithika, M; Prithivikumaran, N; Jeyakumaran, 2014). Then, crystalline phase of synthetized CdO NPs was analyzed by Xray diffraction (XRD), size and morphology were also characterized using HR-TEM (high-resolution transmission electron microscopy) (see AL Naggar et al., 2018). In general, mean crystalline size estimated for CdO NPs was 69.84 nm **(Table S1).**

### 2.2. Fly strain and culture

Wild type *D. melanogaster* flies (Canton-s), obtained from Bloomington *Drosophila* stock center, were used in all experiments. Flies were grown on regular *Drosophila* food media containing cornmeal-agar (14–15 g agar, 18.5 g yeast, 61 g glucose, 30.5 g sucrose, 101 g corn meal/L, then kept at 25 °C, 50-60 % relative humidity (RH) with an 18/6-h light/dark cycle) (El-Kholy et al., 2015).

### 2.3. Determination of LC_50_

Cadmium oxide NPs stock solution (20 mg/ml) was prepared by placing 200 mg of CdO NPs in 10 ml of 10% sucrose solution. CdO NPs solution was then sonicated for 30 min using an ultrasonic system (Powersonic 405) before use. Five serially diluted concentrations of synthetized CdO NPs (0.02, 0.06, 0.18, 0.54, 1.62 mg/ml) in standard medium were prepared for bioassays with adult *D. melanogaster*. For each concentration, 10 ml of CdO NPs spiked medium were poured into glass Petri dishes (7 cm in diameter). Then 3-day old *Drosophila* adults (n=10) were transferred to each Petri dish containing medium spiked with CdO NPs. Exposures to each concentration were performed in triplicate. Preliminary findings of the LC50 bioassay indicated that the data obtained after 24 hours and 48 hours of treatment did not meet the requirements for LC50 determination. Mortality was then recorded daily for 4 days, and the total mortality for each concentration measured was then determined. LC_50_ for CdO NPs was then calculated using the LdP Line^R^ program using the log-probit model (Ehabsoft (http://www.ehabsoft.com/ldpline).

### 2.4. Effects of a sublethal concentration of CdO NPs on life history traits of *D. melanogaster*

The LC_50_ of CdO NPs against adult *D. melanogaster* was 0.17 mg/ml. Then, in order to determine potential adverse effects of exposure to lesser concentrations of CdO NPs on survival, developmental time, fecundity, cellular structure and on detoxification related genes of *D. melanogaster*, flies were chronically exposed for 7 days to a non-treated (control) or a sublethal concentration (0.03 mg CdO NPs/ml, which was 20% of the LC_50_, in supplemented media.

For potential effects on survival, ten pairs of newly hatched adults (F_0_) were placed into vials contained a non-treated (control) or a sublethal concentration of CdO NPs-supplemented media. Then we routinely changed both the control and the treated food media every two days until the experiment ended. Mortality was recorded daily. Dead flies were removed from both control and treated vials and sexes of dead flies determined. Three replicates per treatment were tested.

The developmental time and eclosion rate of progeny (F_1_) generation (egg-adults) of F_0_ flies that were exposed to a sublethal concentration of CdO NPs in supplemented media were recorded and the number of pupae (normal, dark and hatched pupae) was determined in each vial compared to that of F_1_ progeny of F_0_ non-treated (control) flies (Rand et al., 2014). Three replicates per treatment were tested. To check for any long-term phenotypic malformations, adults (3-5 days old) of F_1_ flies (n=30) were checked relative to F_1_ of non-exposed (control) flies by use of an Olympus BX61 microscope.

Potential effects on fecundity were assessed by mating newly emerged, virgin females (n=3) and males (n=5) of F_0_ flies that were exposed for 7 days to a non-treated (control) or a sublethal concentration of CdO NPs in supplemented media. Then, flies were transferred into vials containing food without CdO NPs but supplemented with blue dye. Laid eggs were counted daily for 2 days. Fecundity was calculated as number of eggs laid per female per day. Potential long-term effects of CdO NPs on fecundity of F_1_ flies was also investigated. To do that, virgin females (n=3) and males (n=5) F_1_ flies were also mated and allowed to lay eggs on normal cultural media and the number of eggs laid per female per day have been counted for 2 days compared to control F_1_ females. Three replicates per treatment were tested.

To quantify possible effects of short-term exposure to CdO NPs on both gene expression and mid gut cell structure of *D. melanogaster*, subsamples of F_0_ flies that had been exposed for 7 days to a non-treated (control) or a sublethal concentration of CdO NPs in supplemented media and their progeny F_1_ flies were collected. Ten individuals (five males and five females) per treatment frozen at −80 °C. For ultrastructural investigations, only F_0_ flies were investigated as described below (section 2.4.3). In all experiments only F_0_ flies were exposed for 7 days to a non-treated (control) or a sublethal concentration of CdO NPs in supplemented media.

#### 2.4.1. Climbing assay

To test possible adverse effects of short-term exposure to CdO NPs on F_0_ adult *D. melanogaster* locomotion, an assay based on negative geotaxis was used as described previously (Stephano et al., 2018). Briefly, newly emerged male flies (n=10) were exposed for 7 days to a non-treated (control) or a sublethal concentration (20% of LC_50_) of CdO NPs-supplemented media, then transferred into an empty 100 ml glass cylinder, gently tapped to the bottom. After 10 min acclimation at room temperature, upward movement of controls and treated flies to the top of the cylinder was videotaped for 30 sec. Speed of climbing (cm/sec) for each individual was then calculated from recorded videos using ImageJ software (version 1.2). Potential long-lasting effects of CdO NPs on locomotion of progeny F_1_ larva were also assessed compared to F_1_ larva of control flies using the same assay. Three replicates per treatment were tested.

#### 2.4.2. Gene expression

Real-time, quantitative polymerase chain reactions (RT-qPCR) were used to quantify effects of CdO NPs on expressions of five detoxification genes; four genes encoding for metallothioneins (MTn A-D) and one gene related to glutathione S-transferase (GSTD2) that have been investigated during previous studies (Egli et al., 2006; Yepiskoposyan et al., 2006a). Primers for all genes were given previously (Sawicki et al., 2003; Southon et al., 2004) **(Table S2).** Total RNA was isolated from a composite sample of 10 individual F_0_ or F_1_ flies (five males and five females) by use of an RNA extraction kit (Thermo scientific), then cDNA was synthesized from RNA extracts according to the manufacturer’s protocol (see Al Naggar et al., 2019). Overall, qPCR was performed on RNA isolated from three composite samples (n=10) per treatment. For each target gene, abundance of transcripts was quantified using 2^-ΔΔCT^ method (Livak and Schmittgen, 2001).

#### 2.4.3. Transmission Electron Microscopy (TEM)

Midguts from three composite samples (n= 10) of CdO NPs-treated and untreated F0 flies were used and processed according to the method of (Reynolds, 1963), then studied and photographed using an electron transmission microscope JEM-1200EX (JEOL, Japan) at an accelerating voltage of 60 kV. (for more details, see Dabour et al., 2019)

### 2.2. Statistical analysis

Data were analyzed using GraphPad Prism version 8.00 for Windows (www.graphpad.com). To better estimate the normality and homogeneity of the variance, the data were converted to log10 when necessary. The impact of treatments on survival of F_0_ flies has been assessed by log-rank (Mantel cox) paired test, *p* < 0.0083 after Bonferroni correction. Differences in developmental time and eclosion rate were assessed by Student’s *t*-test. Differences in climbing ability of F_0_ and F_1_ larva were assessed by Mann Whitney test. Effect of treatment on fecundity of F_0_ and F1 flies were assessed by Two-way RM ANOVA. Effects on gene expression between treatments were analyzed by one-way ANOVA followed by Tukey’s post hoc test. An alpha level of 0.05 was used to define significance for all tests.

## 3. Results

### 3.1. Effects on life history traits

Chronic exposure of F_0_ *D. melanogaster* flies to food media spiked with a sublethal concentration of CdO NPs for 7 days, significantly reduced survival of both male and female flies as compared to controls (log-rank (Mantel cox) paired test, *p* < 0.0001, after Bonferroni correction). All treated flies died after 17 days. Survival of males did not differ significantly compared to females regardless of whether they were exposed to CdO NPs or not (log-rank (Mantel cox) paired test, *p* > 0.05) **(Fig.1a).**

**Fig 1.**
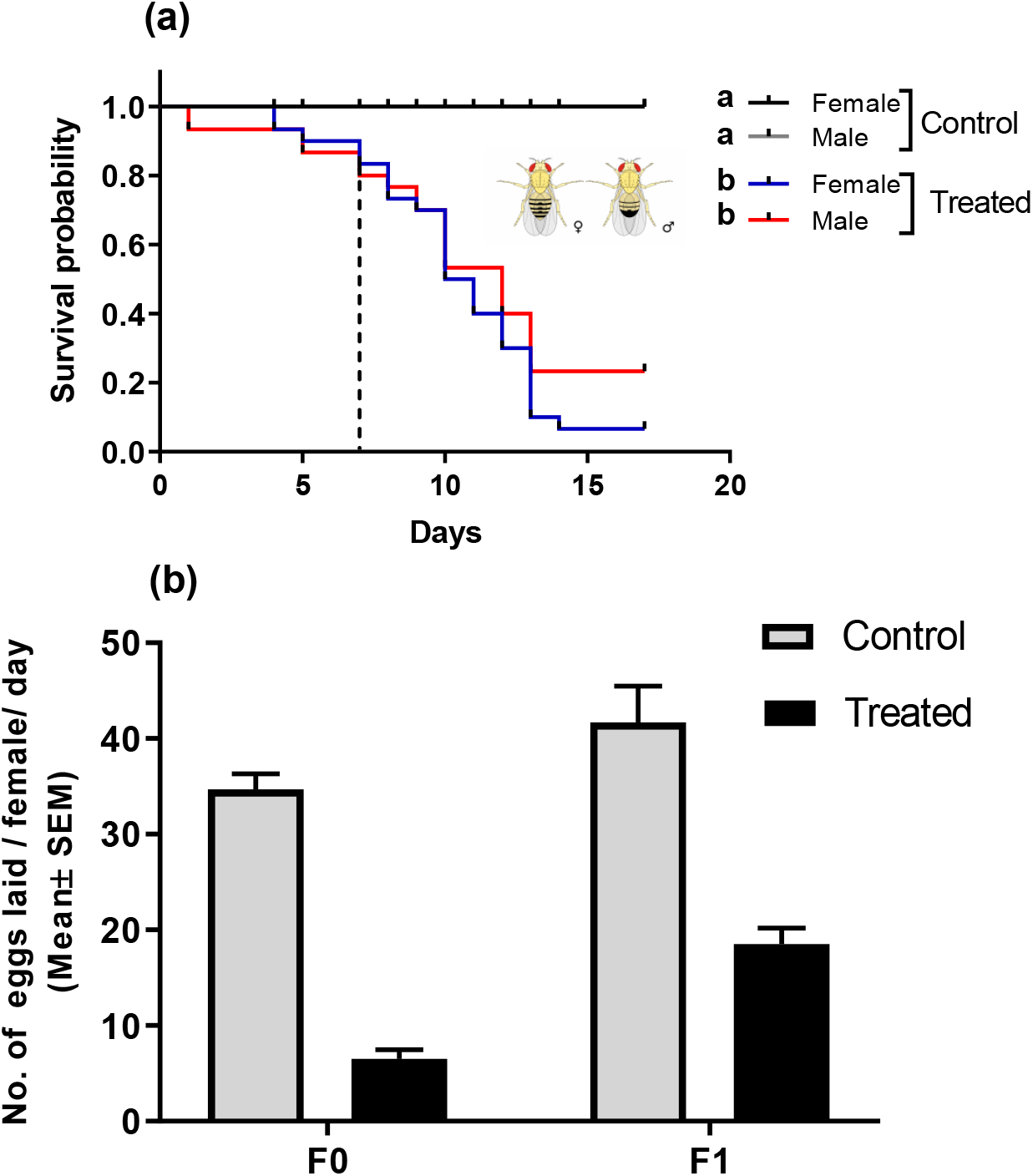
Kaplan–Meier plot showing effects of CdO NPs on survival of male and female F_0_ *D. melanogaster flies* (a) and fecundity (b). **(a)** Adult flies were exposed for 7 days to a control or food media spiked with a sublethal concentration (0.03 mg/ml) of CdO NPs and then transferred to normal cultural media every two days. Different lowercase letters indicate statistical differences between treatments after Bonferroni correction (log-rank (Mantel cox) paired test, *p* < 0.0001). Dashed line indicates the end of CdO NPs exposure time. (b) Number of eggs laid/female/day (mean ± sem) of F_0_ flies (parents) that exposed to a control or food media spiked with a sublethal concentration (0.03 mg/ml) of CdO NPs for 7 days and their F1 progeny. Number of eggs laid were significantly different between both treated and non-treated flies and also between F_0_ and F_1_ (Two-way RM ANOVA, *p* < 0.001).

Developmental time and rates of eclosion of *D. melanogaster* F_1_ generation (egg-adults) of F_0_ flies that were exposed to lesser concentrations of CdO NPs resulted in adverse effects compared to controls **(Table 1)**. Eggs hatched after one day exposure in both exposed and non-exposed control groups. The time required for larval and pupal development was longer in CdO NPs-treated flies, however this difference was not statistically significant compared to the control (Student’s *t*-test*, P* > 0.05). Developmental time (egg-adult) was significantly longer in flies exposed to CdO NPs (11.72 ± 0.60 days) compared with the control (8 ± 0.00 days) (*P* < 0.001). Eclosion rate (%) was also significantly less in treated flies (12.38 % ± 2.94) compared to control (69.50 % ± 0.60) (*P* < 0.001) **(Table 1).**

**Table 1.** Developmental time in days and percentage of pupa (Mean ± SD) of *Drosophila melanogaster* that exposed to a control or food media spiked with a sublethal concentration (0.03 mg/ml) of CdO NPs for 7 days.

| Treatment | Development time (days) |  |  |  | Eclosion assay (%) |  |  |
| --- | --- | --- | --- | --- | --- | --- | --- |
|  | Egg deposition | Larvae | Pupae | Adult | Normal pupa | Black pupa | Hatched pupa |
| Control | 1 ± 0.00 <sup>A</sup> | 2 ± 0.00 <sup>A</sup> | 5 ± 0.00 <sup>A</sup> | 8 ± 0.00 <sup>A</sup> | 17.78 ± 1.47 <sup>A</sup> | 12.72 ± 1.10 <sup>A</sup> | 69.50 ± 0.60 <sup>A</sup> |
| CdO NPs | 1 ± 0.00 <sup>A</sup> | 3 ± 0.00 <sup>A</sup> | 7 ± 0.00 <sup>A</sup> | 11.72 ± 0.60 <sup>B</sup> | 72.74 ± 4.22 <sup>B</sup> | 14.88 ± 1.30 <sup>A</sup> | 12.38 ± 2.94 <sup>B</sup> |

\*Different uppercase letters denote significant differences in development time (days) for each stage and the different number of each pupa type compared with the control (Student' s *t*-test, *P* <0.001).

Fecundity of F_0_ females of *D. melanogaster* that exposed for 7 days to a control or food media spiked with a sublethal concentration of CdO NPs and their F_1_ progeny is shown in **Figure 1 (b)**. Number of eggs laid per female per day were significantly different between both treated and non-treated flies and also between F_0_ and F_1_ flies (Two-way RM ANOVA, *p* < 0.001).

### 3.2. Effects on climbing ability

Short-term exposure of male F_0_ flies for 7 days to a sublethal concentration of CdO NPs affected their instinctive negative geotaxis behavior. Exposed flies were excessively disturbed, hyperactive preferred to jump or use their wings, therefore, climbed in short paths. Only data of flies walked up vertically against the gravity are considered. Their climbing speed significantly impaired compared to control (Student’ s *t*-test, *P* < 0.001) **(Fig. 2a).** There was no significant difference in climbing ability of F_1_ larva of treated flies compared to F_1_ larva of controls (Student’ s *t*-test, *P* > 0.05) **(Fig. 2b).**

**Fig.2.**
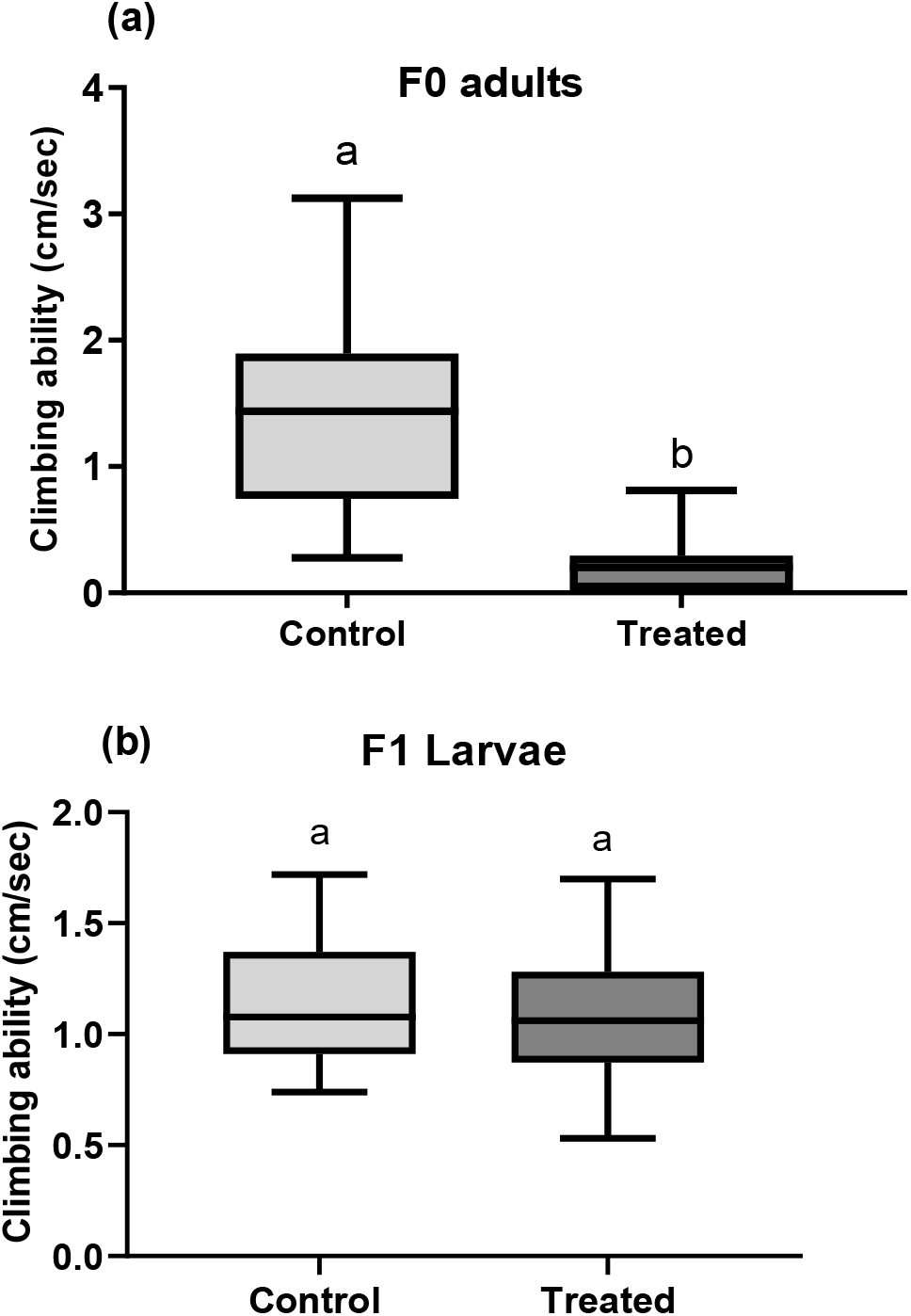
Climbing ability (cm/sec) of (a) F_0_ males (parents) that were exposed to a control or food media spiked with sublethal concentrations of CdO NPs for 7 days and (b) their progeny F_1_ larva. Symbols on the box plot represent maximum and minimum values (n=32) (whiskers: ┬ ┴), mean values (-). Different letters denote significant difference from the control (Mann Whitney test, *P* < 0.0001).

### 3.3. Effects on gene expression

Abundance of transcripts of genes coding for metallothionein (MTn A-D) and glutathione S-transferase (GSTD2) involved in detoxification of metals in both *D. melanogaster* F_0_ and F_1_ flies are shown in Figure 3. Genes encoding for metallothionein (MTn A-D) were significantly up-regulated in Cd) NPs-exposed F_0_ flies and their progeny F_1_ flies compared to F_0_ and F_1_ of controls (*p* < 0.05) **(Fig. 3).** Additionally, expressions of MTn A-C encoding genes were greater in F_1_ flies compared to F_0_ flies. Although expression of GSTD2 was significantly upregulated in F_0_ and F_1_ flies, this difference between flies exposed to CdO NPs and controls was not statistically significant (*p* > 0.05) **(Fig. 3).**

**Fig.3.**
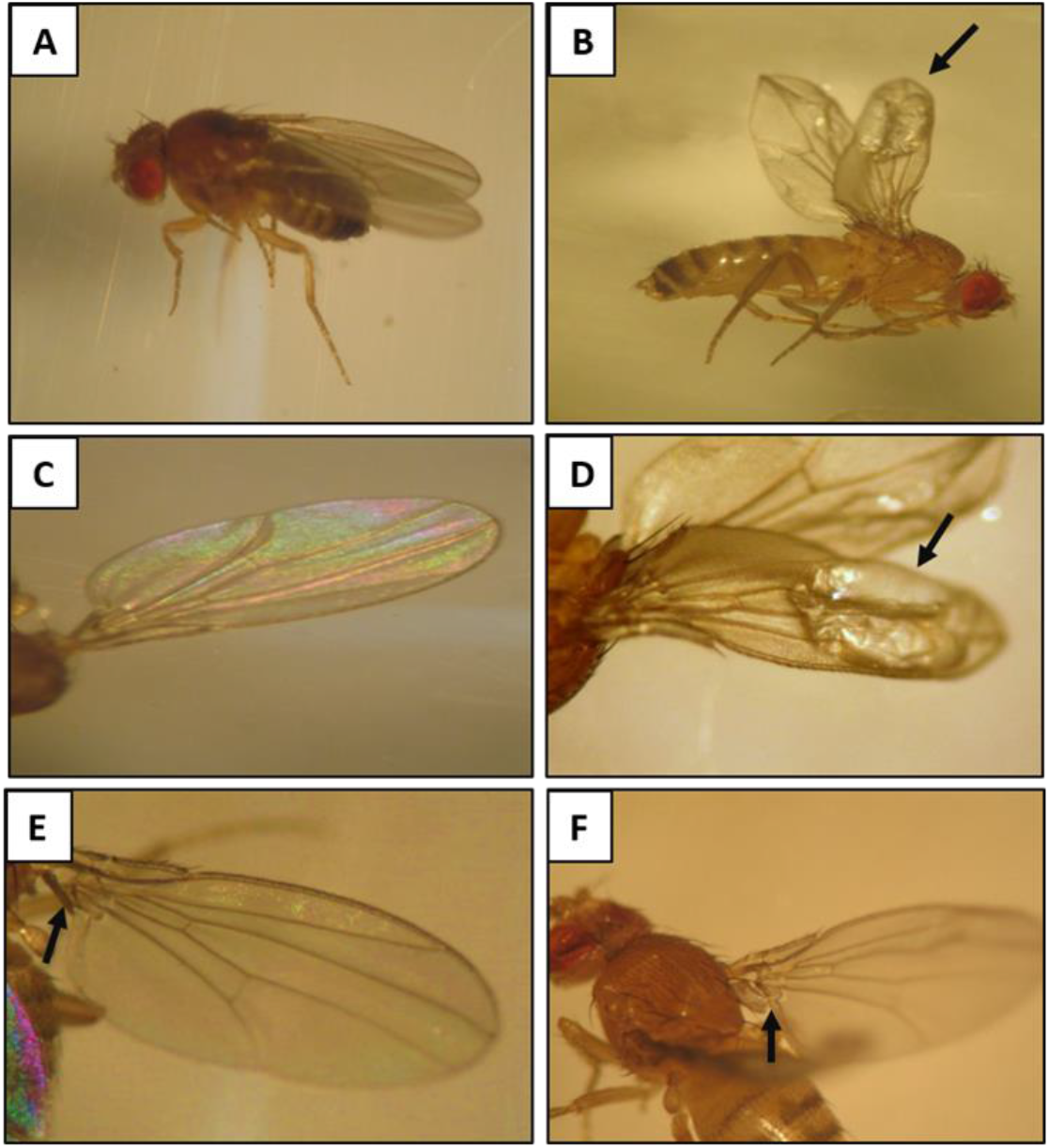
Fold-change in abundances of transcripts of metallothionein (MTn A-D) and glutathione S-transferase encoding genes involved in detoxification of heavy metals in *Drosophila melanogaster* F_0_ and their F_1_ progeny flies. Adult F_0_ flies (parents) have been exposed to a control or food media spiked with a sublethal concentration (0.03 mg/ml) of CdO NPs for 7 days. Bars represent the mean ± SEM concentration of three samples. Different lowercase letters denote significant differences among treatments (one-way analysis of variance with Tukey’s post-hoc test, *P* <0.05).

### 3.4. Phenotypic effects and ultrastructure observations by TEM

F1 flies (n=30) of F_0_ that exposed for a short-term to a sublethal concentration of CdO NPs were carefully checked for any physical abnormalities in body size and appendages. Malformations of wings of 83.3 % (n=25) were observed and flies exhibited blisters and bubbles at the apical area of the wings and reduction of the axillary cord at the jugal area of the wings compared to control **(Fig. 4.)**.

**Fig.4.**
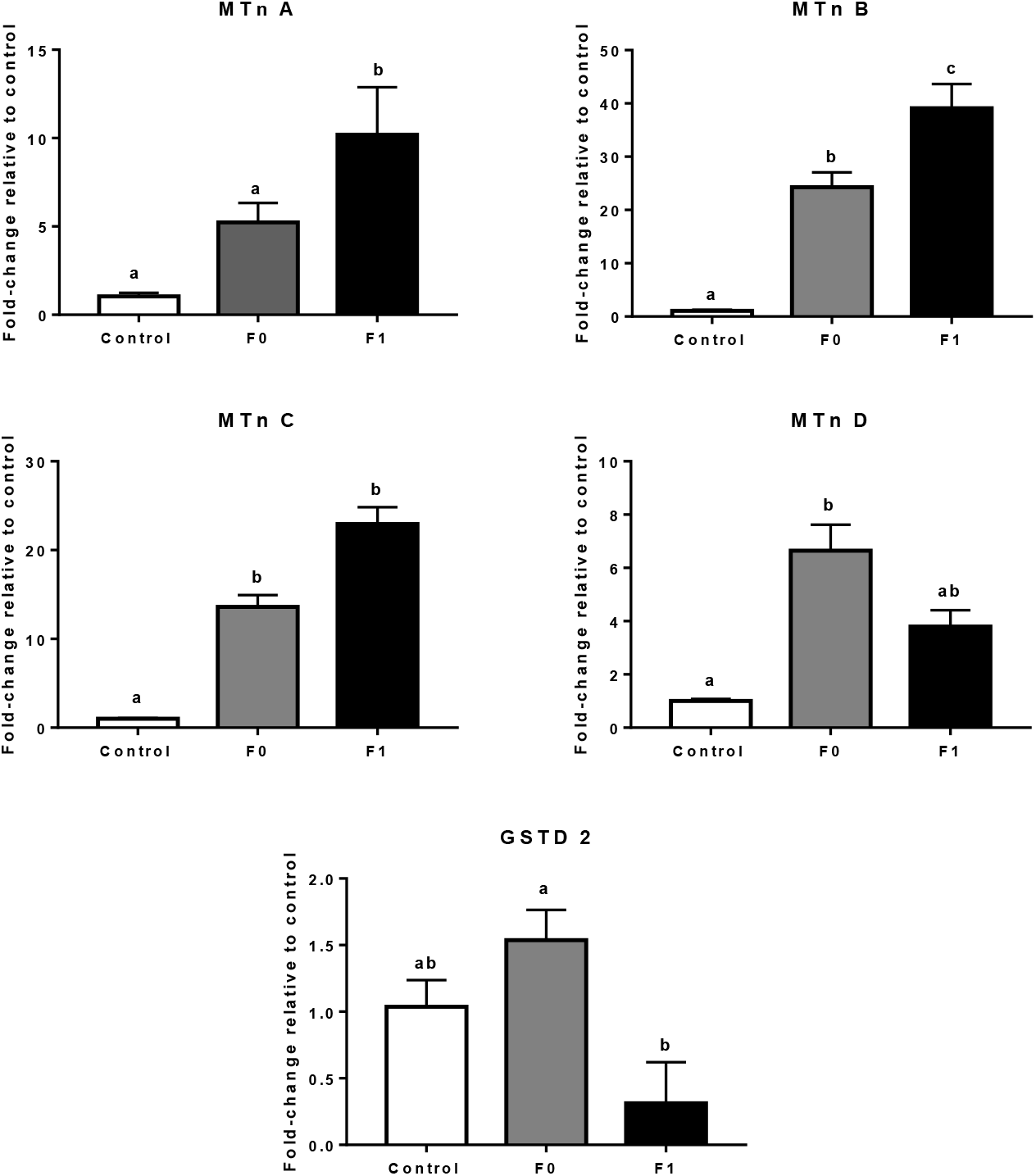
Phenotypic effects of CdO NPs on wings of 3-days old adults F_1_ *Drosophila melanogaster* flies compared to F_1_ control flies. Parents (F_0_) of these flies have been exposed for 7 days to a control or food media spiked with a sublethal concentration (0.03 mg/ml) of CdO NPs. A, C and E are F_1_ of control flies showing normal wings. B, D, F are F_1_ of treated flies showing malformation in wings. Note, the blisters & bubbles at the apical margin of the wings and reduction of the axillary cord at the jugal area of the wings (black arrows).

Bioaccumulation of CdO NPs in midgut cells of F_0_ flies that were exposed for 7 days to food media spiked with a sublethal concentration of CdO NPs is shown in **Figure S1**. Cellular alterations in mid gut cells of F_0_ treated flies as compared to control flies are shown in **Figure 5**. Midgut cells of non-treated (control) F_0_ flies exhibited typical columnar cell morphology with the apical border, which was straight bearing multiple, long filaments like microvilli. Large and dense mitochondria, rER and an oval nucleus were also found **(Fig. 5 A, C, E)**. Ultrastructure’s of midgut epithelial cells in flies exposed to CdO NPs were adversely affected. Lysis of the smooth endoplasmic reticulum(sER), microvilli were found to be fragmented and large lytic region observed. Some mitochondria have been found swollen, had matrix lysis, mitochondrial crystal breakage and autophagosomal appearance. Moreover, we also observed lysis of the nuclear membrane and in rER membranes, its layered structure was thus lost **(Fig. 5 B, D, F)**.

**Fig 5.**
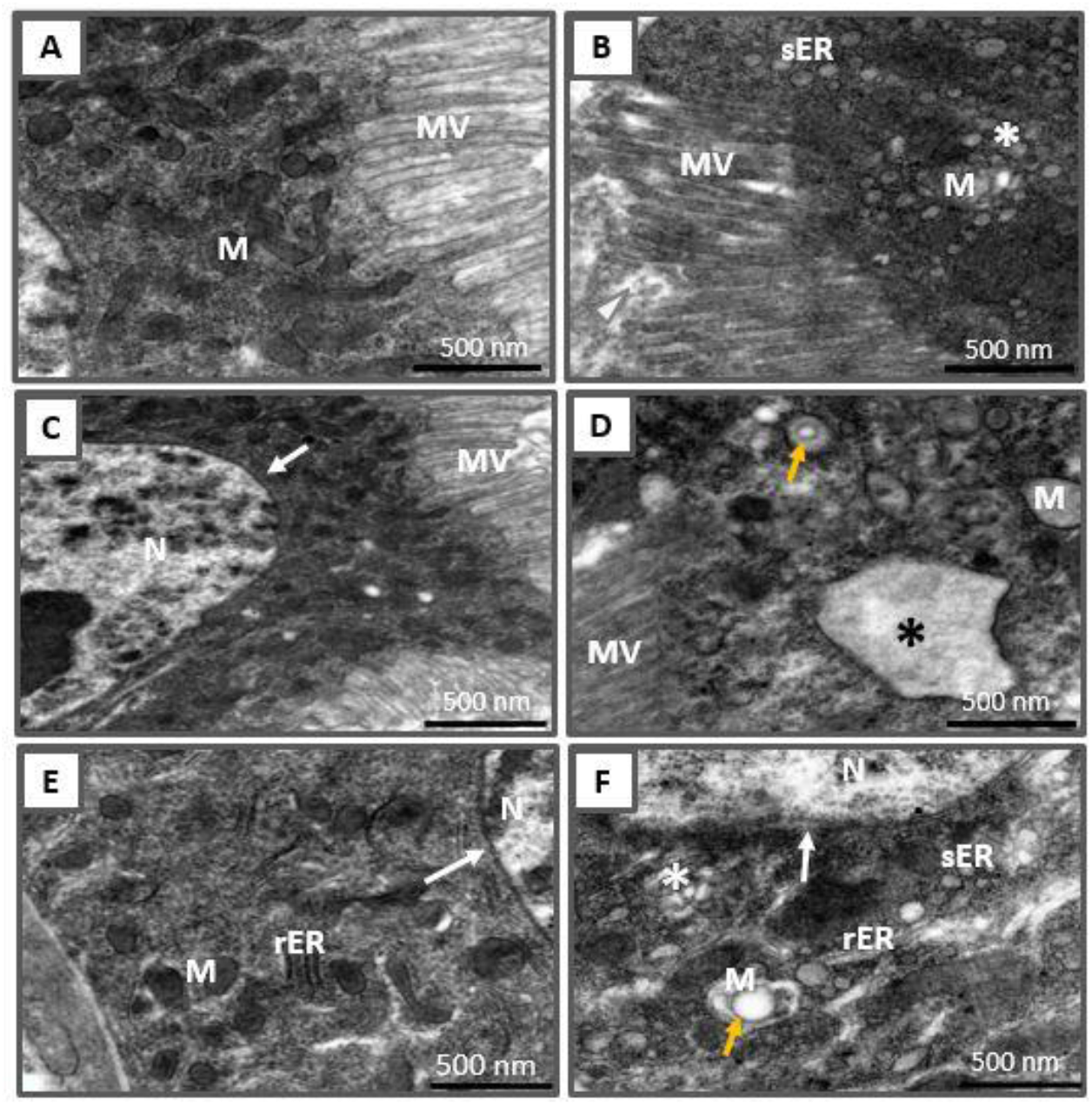
Transmission electron microscopy photomicrographs of midgut cells of F_0_ *Drosophila melanogaster* flies that exposed to a sublethal concentration (20 % of LC_50_) of CdO NPs for 7 days compared to control. A, C and E Control group, exhibiting typical morphology of columnar cells with the apical border that was straight bearing numerous, long filament like microvilli. Note, columnar cells with oval nucleus, abundant and dense mitochondria and rough endoplasmic reticulum (rER). B, D and F flies exposed to a sublethal concentration of CdO NPs (0.03 mg ml^-1^). Note, lysis of smooth endoplasmic reticulum (sER) (white stars), microvilli appeared fragmented (white arrowhead) and large lytic area (black star). Some mitochondria were found swollen and showed lysis of matrix and breakage of mitochondrial cristae and appeared autophagosome (burlywood arrows). Note, the lysis in the nuclear membrane (white arrow) and in rER. N Nuclei; sER smooth endoplasmic reticulum; rER rough endoplasmic reticulum MV microvilli; M mitochondria.

## 4. Discussion

Exposure of aquatic and terrestrial organisms to potentially toxic metals and metalloids can have adverse effects (Ali et al., 2019). Even at sublethal concentrations, metals and metalloids can cause toxic effects (AL Naggar et al., 2018; Dabour et al., 2019). Given a substantial amount of existing literature on potential risks associated with exposure to emerging nanomaterials (Exbrayat et al., 2015), to our knowledge, this is the first study to investigate and report adverse long lasting effects on key life history traits of *D. melanogaster* well beyond exposure times to CdO NPs even at a lesser concentration.

The results obtained showed that CdO NPs were more toxic “exert adverse effects” to *D. melanogaster* flies than were cadmium chloride (CdCl2) soluble salts. Where, the calculated median lethal concentration (LC_50_) of CdO NPs was (0.17 mg/ml), while the LC_50_ values of CdCl2 ranged from 0.30 to 0.51 mg/ml according to various fly genotypes (Christie et al., 1985). These results are in agreement with the hypothesis that NPs derived from Cd and possibly other metal oxides will be more potent than their metal salts because of their specific properties (Sukhanova et al., 2018).

In the current study, short-term exposure of adult *D. melanogaster* to a sublethal concentration of CdO NPs adversely affected all studied life history traits of the fly including survival, developmental time and reproductive fitness. Such results are in line with previous studies revealing that exposure to Cd, either as salts (CdCl2) or as nano-sized (CdO) can cause toxic effects in *D. melanogaster* as well as organisms (AL Naggar et al., 2018; Bixler and Schnee, 2018; Blum et al., 2012; Dabour et al., 2019; Hu et al., 2019; Mathew and NB, 2018). Inhalation of CdO NPs during pregnancy negatively affects fecundity and inhibits the development of fetal and postnatal offspring of mice (Blum et al., 2012). Here, the flies exposed to CdO NPs by oral intake and long-lasting effects beyond direct exposure times were further investigated by assessing the fecundity of both parents (F_0_) flies and its non-treated progeny (F_1_) flies where, the fecundity of F_1_ flies were significantly reduced (−50%) compared with fecundity of unexposed F_1_ flies. These findings also match the results of previous studies, which indicated that the presence of Cd in the environment may impair the fitness of *D. melanogaster* adulthood, even at lesser concentrations for short durations (Bixler and Schnee, 2018; Mathew and NB, 2018), because it might negatively affect expressions of genes associated with reproduction of *D. melanogaster* and trigger the transcription of defense-related genes (Hu et al., 2019).

Irritable and fast climbs point to a chronic neuronal motor defect in *D. melanogaster* flies (Anand et al., 2019). Here, climbing behavior was impaired in only F_0_ flies exposed to CdO NPs. In an earlier study, in which honey bees were exposed to a sublethal concentration of CdO NPs, bees also showed malaise-like’ behaviors (AL Naggar et al., 2018). These findings confirmed the neurotoxicity of CdO NPs and future studies are therefore required to explore underlying mechanisms of its neurotoxicity on living organisms.

Normally, wings of insects are smooth, consisting of single dorsal and ventral epithelial layers kept together by cell adhesion and any interaction with cell adhesion leads to the apposition of wing epithelial sheets and thus to an aberrant 2D wing structure (Bilousov et al., 2014). Here, chronic exposure of F_0_ flies led to blistered wings in their F_1_ progeny flies compared to F_1_ control flies. Similarly, adverse birth outcomes were also detected however, as a result of moderate prenatal Cd exposure of pregnant women (Taylor et al., 2016). Birth defects detected due to either dietary CdO NPs exposure in the current study or due to inhaled CdO NPs (Blum et al., 2012) emphasize the great toxicity of nano-sized CdO even at sublethal levels and give a clear public health message to women who are pregnant and those of childbearing age who are exposed to CdO NPs at work.

An intrinsic characteristic of contaminated species is their ability to adjust to such toxin concentrations by switching to a variety of countervailing detoxification mechanisms or modifying the expression and function of several enzymes. Metallothionin (Mtn) is a ubiquitous, lightweight, cysteine-rich protein capable of binding essential heavy metal cations, such as Zn and Cu, or heavy metals with no known biological function such as Hg, Cd, Ag, etc. (Yepiskoposyan et al., 2006b). In the present study, MTn (A-D) encoding genes were significantly induced by CdO NPs in both F_0_ and F_1_ flies compared to control. This is noteworthy because there was a two-week time interval between the exposure of F_0_ parent flies to CdO NPs and the measures of gene expression in their F_1_ progeny flies, which is almost spanning the developmental time (egg-adult) of the fly (Perveen, 2018). This was in addition to the phenotype effects observed in wings of F_1_ flies indicates the long-term adverse effects of CdO NPs even at low exposure levels and for short periods during early adulthood.

Cadmium is a non-metal that can be deposited in animal tissues, especially if it is found in nanosized materials capable of disrupting physiological functions that cause significant internal tissue damage (Suganya et al., 2016). Common cytological changes observed in epithelial cells of F_0_ flies in the current study were swelling and lysis of both mitochondria and sER and lysis of nuclear and rER membranes. Moreover, microvilli appeared fragmented and large lytic area were observed as well. Such ultrastructural changes reflect the key features of cell necrosis and apoptosis (Elmore, 2007; Proskuryakov et al., 2003) and are comparable to those found in the midgut of the honey bees exposed to CdO NPs (Dabour et al., 2019). Moreover, it could explain the observed long lasting adverse effects on studied life history traits of *D. melanogaster*.

## 5. Conclusions

Extensive production and use of metal (oxide) nanoparticles increase the potential for their release into the environment. Results of this study showed for the first time that short-term exposure to a sublethal concentration of CdO NPs is sufficient to cause long-lasting, harmful effects on life history traits of the fruit fly, which might also occur in other organisms. Deformities in wings of F_1_ progeny flies were also observed. Common cytological changes in epithelial cells were mitochondrial swelling and lysis, and lysis of nuclear membrane and rER membranes that were nearly identical to those seen in other insects or invertebrates exposed to these metals. Taken together, the findings of this study provide clear insight into CdO NP’s possible danger to living organisms using *D. melanogaster* as an *in vivo* model. Nanotechnologies based on cadmium (Cd) pose risks to humans and the environment and their use needs to be regulated and the environment monitored for potential exposure.

## 6. Acknowledgments

This study was partly funded by grants for scientific research from the Faculty Science, Tanta University, Egypt.

## 7. Supplementary material

**Table S1.** XRD data of CdO nanoparticles prepared at 0.5 M.

**Table S2.** List of primers used for quantification of abundances of transcripts in *Drosophila melanogaster* by RT-qPCR

**Figure S1.** Transmission electron microscopy photomicrographs of mid gut cells of F_0_ flies that were exposed for 7 days to food media spiked with a sublethal concentration of CdO NPs showing the internalization and bioaccumulation and of CdO NPs.

## Notes

### Competing Interest Statement

The authors have declared no competing interest.

