## Supplementary material for "Consequences of a short-term exposure to a sub lethal concentration of CdO nanoparticles on key life history traits in the fruit fly (*Drosophila melanogaster*)": Supporting Information.docx

* Corresponding author:

Yahya Al Naggar, Ph.D.

**Table S1.** XRD data of CdO nanoparticles prepared at 0.5 M.

| **Metal** | **d-spacing**  **(Å) Observed** | **Intensity**  **(cps)** | **FWHM**  **β (deg)** | **(2Theta)** | **Grain size**  **(nm)** |
| --- | --- | --- | --- | --- | --- |
| **CdO** | 2.71 | 100.00 | 0.17 | 33.03 | 50.93 |
|  | 2.34 | 91.25 | 0.17 | 38.34 | 51.70 |
|  | 1.66 | 55.26 | 0.10 | 55.32 | 95.64 |
|  | 1.41 | 34.97 | 0.12 | 65.94 | 81.11 |

**Table S2.** List of primers used for quantification of abundances of transcripts in *Drosophila melanogaster* by RT-qPCR

| **Locus** | | | **Category** | | **Sequences** | | **References** |
| --- | --- | --- | --- | --- | --- | --- | --- |
| **Metallothionein (MtnA)** | | Detoxification | | | F: CCTGCAACTGCGGATCT | | (Southon et al., (2004) |
|  |  |  |  |  | R: CGCAGGCGGATTTCTT | |  |
| **MtnB** | | Detoxification | | | F: ATGGTTTGCAAGGGTTGTG | | (Southon et al., (2004) |
|  |  |  |  |  | R: TTGCAGGCGCAGTTGT | |  |
| **MtnC** | | Detoxification | | | F: GCACTTGCAGTCCTGATTACAG | | (Southon et al., (2004) |
|  |  |  |  |  | R: GCACTTGCAGTCCTGATTACAG | |  |
| **MtnD** | | Detoxification | | | F: AGTGCTCCGCCACCAA | | (Southon et al., (2004) |
|  |  |  |  |  | R: TGTCCTTGGGTCCGTTCT | |  |
| **Glutathione S transferase (GSTD2)** | | Detoxification | | | F:GCAATATCCCCCATATGGACTTTTACTAC | | (Sawicki et al., 2003) |
|  |  |  |  |  | R: ATAGTTGGGATCCCGATCACTTAGC | | (Li et al., 2015) |
| **RpL32** | | House keeping | | | F: GCTAAGCTGTCGCACAAATG | |  |
|  |  |  |  |  | R: GTTCGATCCGTAACCGATGT | |  |

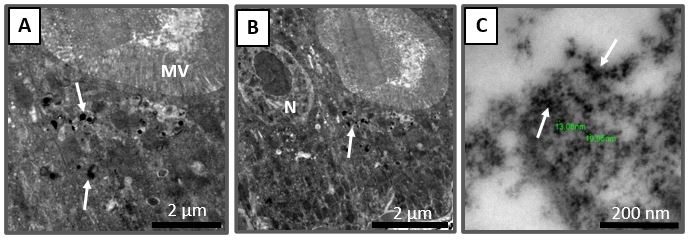

**Figure S1.** Transmission electron microscopy photomicrographs of mid gut cells of F_0_ flies that were exposed for 7 days to food media spiked with a sublethal concentration of CdO NPs showing the internalization and bioaccumulation and of CdO NPs (white arrows).

.
